# Specific decreasing of Na^+^ channel expression on the lateral membrane of cardiomyocytes causes fatal arrhythmias in Brugada syndrome

**DOI:** 10.1101/805986

**Authors:** Kunichika Tsumoto, Takashi Ashihara, Narumi Naito, Takao Shimamoto, Akira Amano, Yasutaka Kurata, Yoshihisa Kurachi

## Abstract

One phenotypic feature of Brugada syndrome (BrS) is slowed conduction due to the reduction (loss-of-function) of Na^+^ channels. In contrast, recent clinical observations in BrS patients highlighted the poor correlation between the phenotype (typical ECG change or lethal arrhythmia) and the genotype (*SCN5A* mutation). Inspired by our previous theoretical study which showed that reduced Na^+^ channels in the lateral membrane (LM) of ventricular myocytes caused the slowing of conduction under myocardial ischemia, we hypothesized that a loss-of-function of Na^+^ channels caused by the decreases in Na^+^ channel expression within myocytes leads to phase-2 reentry (P2R), the major triggering mechanism of lethal arrhythmias in BrS. We constructed an *in silico* human ventricular myocardial strand and ring models, and investigated the relationship between the subcellular Na^+^ channel distribution and P2R. Reducing Na^+^ channel expression in the LM of each myocyte caused not only the notch-and-dome but also loss-of-dome type action potentials and slowed conduction, both of which are typically observed in BrS patients. Furthermore, we showed that both the reduction in Na^+^ channels on the LM of each myocyte and tissue-level heterogeneity of Na^+^ channel expression were essential for P2R as well as P2R-mediated reentrant excitation. Our data suggest that the alteration in subcellular Na^+^ channel distribution together with a tissue-level heterogeneity of Na^+^ channels can cause arrhythmogenesis in BrS.

## Introduction

Cardiac Na^+^ channel current (*I*_Na_) that controls the excitability of cardiomyocytes mainly contributes to the action potential (AP) initiation and its propagation. Na^+^ channel dysfunction in congenital or acquired heart diseases leads to decreases in the *I*_Na_, which has been associated with slowed or blocked conduction, resulting in a potentially proarrhythmic substrate. Genetic abnormalities causing Na^+^ channel dysfunction have been linked to many arrhythmogenic diseases (*1*), including sick sinus syndrome, progressive cardiac conduction disorders, and Brugada syndrome (BrS). In contrast, a previously published study suggested that variants of *SCN5A*, which is one of the important disease genes for BrS (*2*), were not necessarily associated with the BrS phenotype (*3*). The loss-of-function mutations in *SCN5A* are not a determinant factor of BrS.

BrS is characterized by the ECGs with both a right bundle-branch block pattern and ST-segment elevation in the right precordial leads (V_1_-V_3_) and by the higher incidence of sudden cardiac death due to ventricular tachycardia/fibrillation (VT/VF) (*4, 5*). It is commonly believed that VT/VF is elicited by the closely coupled premature ventricular contractions via the phase-2 reentry (P2R) mechanism (*6*). This mechanism is caused by an electrotonic current flowing from the region where AP exhibits notch-and-dome morphology to the region where AP shows loss of the phase-2 dome.

Although at first, it has been predicted that a global difference in AP duration (APD) causing a transmural voltage gradient in the right ventricle (RV) is a cause of P2R development (*4*), several experimental studies using an optical mapping system showed that the epicardial local APD difference was involved in the occurrence of P2R (*7, 8*). Thus, the P2R mechanism in BrS is still controversial.

Previous experimental studies (*9, 10*) have produced knock-in mice lacking the SIV (Serine-Isoleucine-Valine) domain (ΔSIV) of the Na_V_1.5 PDZ domain–binding motif that interacts with PDZ proteins (syntrophin and SAP97). Experiments on these mice showed that both Na^+^ channel expression and *I*_Na_ on the lateral membrane (LM), but not on the intercalated discs (IDs), of cardiomyocytes were decreased. This led to slowed conduction at the tissue level. Notably, Shy et al. (*10*) identified a missense mutation within the SIV motif (p.V2016M) from a patient with BrS, suggesting that the local alteration in subcellular Na^+^ channel expression as observed in the ΔSIV mice might be involved in BrS. In a previous theoretical study (*11*), we showed that a subcellular alterations in Na^+^ channel expression induced by myocardial ischemia, specifically reduced number of Na^+^ channels in the LM of cardiomyocytes, was strongly associated with both the decrease in local *I*_Na_ and slowed conduction.

Furthermore, the local decreasing of *I*_Na_ in cardiomyocytes evoked P2R. Therefore, we hypothesized that a local alteration in subcellular Na^+^ channel expression may lead to the development of P2R and VT/VF in BrS. To rigorously examine this hypothesis, we require an analyzing system in which Na^+^ channel expression within each myocyte can be selectively controlled. We, therefore, developed *in silico* human ventricular myocardial strand and ring models reflecting the recent experimental data (*10*) on subcellular Na^+^ channel expression changes observed in the ΔSIV mice. Then, we performed simulations of AP propagation in the *in silico* models and investigated the relationship between the subcellular alterations in Na^+^ channel expression and the occurrence of P2R leading to reentrant arrhythmia onsets in BrS.

## Methods

### Myocardial strand and ring models and subcellular Na^+^ channel distribution

To perform computer simulations examining effects of alterations in subcellular Na^+^ channel distribution on AP propagation behaviors, we used myocardial strand (Fig. 1A*a* and fig. S1) and ring (Fig. 1B) models comprised of cylindrical human ventricular myocytes, each 150 μm in length and 20 μm in diameter (*12*). As shown in Fig. 1A*b*, adjacent myocytes were electrically coupled with both gap junctions and an electric field mechanism (ephaptic coupling) (*13, 14*), the latter of which is an interference effect caused at intercellular cleft space; electrical communication between myocytes through the electric field mechanism is mediated by large negative changes in the extracellular potential elicited within the narrow intercellular cleft space facing the IDs.

**Fig. 1.**
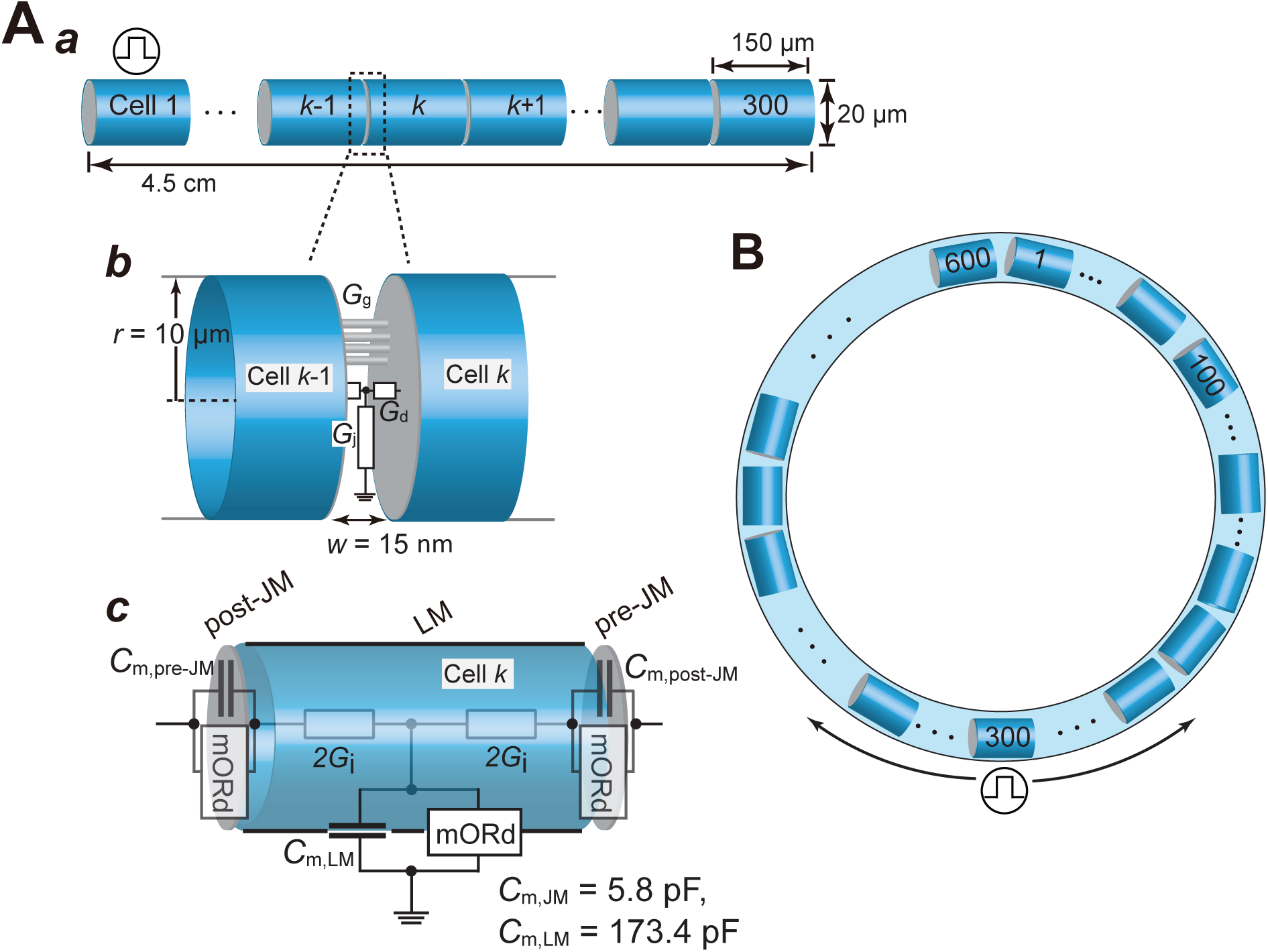
Myocardial strand and ring models. **A** and **B**, Schematic representation of a myocardial strand model comprising cylindrical 300 cells (*a*) and of intercellular junction between myocytes (*b*). The cell membrane in each myocyte is divided into post-junctional, lateral, and pre-junctional segments, and electrophysiological properties of the each membrane segment are represented by a modified O’Hara-Rudy dynamic (mORd) ventricular myocyte model (*c*) (*16, 17*). **B**, Schematic representations of the myocardial ring model comprising 600 cells. Cell #1 (A) and #300 (B) were electrically stimulated at 1 Hz. *G*_g_, gap junction conductance; *G*_j_, radial conductance of intercellular cleft; *G*_d_, axial conductance of intercellular cleft.

As in our previous study (*11*), to achieve the inhomogeneous Na^+^ channel distribution within a myocyte found in humans and other mammals (*15*), the whole cell membrane of each myocyte was divided into three segments (Fig. 1A*c*), one segment for the LM and the other two segments for the junctional membranes (JMs), namely the IDs. Allocating Na^+^ channel conductances to each membrane segment and changing those individually, we altered the subcellular expression of Na^+^ channels. The electrophysiological property of each membrane segment was represented by the O’Hara-Rudy dynamic (ORd) model (*16, 17*), which is the most sophisticated human ventricular AP model to date that extensively validated against experimental data from more than 100 non-diseased human hearts. For better correspondence with the experimental conduction velocity of human RV epicardium (*18*), the formula of fast *I*_Na_ in the ORd model was replaced with that of the ten Tusscher-Panfilov (TP) model (*19*). We set the control conductance of fast *I*_Na_ to 11 nS/pF for the LM (*G*_NaF,LM_) and 44 nS/pF for the JMs (*G*_NaF,JM_) so that *I*_Na_ amplitude obtained from the simulation matched the one recorded experimentally for basal condition (*10*); the conductance of late I_Na_ was also set to 0.0045 nS/pF for the LM (*G*_NaL,LM_) and 0.018 nS/pF for the JMs (*G*_NaL,JM_) as the control condition. Throughout the article, we express the fast and late Na^+^ channel conductances of the JM and LM as percentages of the *G*_NaF,JM_ (*G*_NaL,JM_) and *G*_NaF,LM_ (*G*_NaL,LM_), i.e., %*g*_Na,JM_ and %*g*_Na,LM_, respectively. Specific parameters in the myocardial strand and ring models consisting of the ORd model can be found in the supplemental table S1.

**Table 1.** Relationships of the Na^+^ channel density at the lateral membrane, the regional Na^+^ channel current, and conduction velocity

|  | **100%$g_{\text{Na,L,M}}$** | **35%$g_{\text{Na,L,M}}$** | **7%$g_{\text{Na,L,M}}$** | **0%$g_{\text{Na,L,M}}$** |
| --- | --- | --- | --- | --- |
| Peak $I_{\text{Na,L,M}}$ (pA/pF) | 329.3 | 138.7 | 24.6 | 0.0 |
| CV (cm/s) | 71.4 | 53.6 | 33.3 | 25.4 |

$\%g_{\text{Na,L,M}}$ , a percent $g_{\text{Na}}$ in the lateral membrane (LM) segment of each cell of the strand model; Peak $I_{\text{Na,L,M}}$ , peak value of $I_{\text{Na}}$ across the LM segment of the 150th cell in the myocardial strand model; CV, conduction velocity.

### Simulations

Numerical calculations were performed as described previously (*11*) and details were provided in Supplementary Materials. Pacing stimuli at threefold diastolic threshold with a basic cycle length of 1 s were applied to a myocyte located at the end of the myocardial strand (the 300th myocyte in the myocardial ring) and repeated 30 times to minimize transient responses in each simulation. The membrane potential was calculated using the forward Euler method with a 1 μs time step to compensate for the reduced size of the discretized space and for the acute change in the membrane potential at the IDs due to the electric field mechanism.

To test whether the sympathetic activity inhibits the P2R development, we additionally performed simulations of AP propagation using the myocardial strand model under the condition of *β*-adrenergic stimulation (*β*-AS) mimicking sympathetic activation. Modifications of parameters for simulating the condition of *β*-AS (*20, 21*) were listed in the supplemental table S2.

**Table 2.**
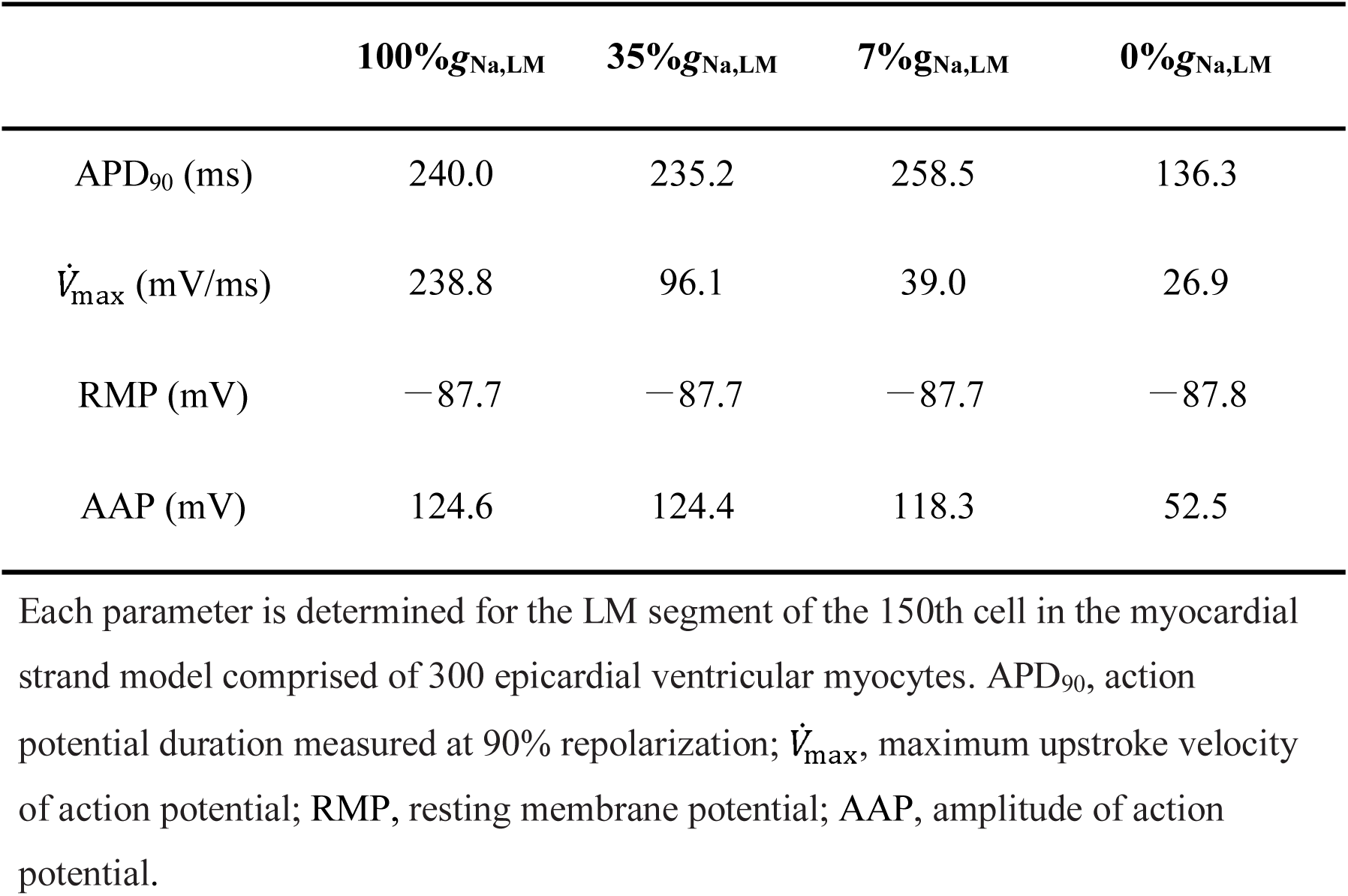
Effects of changes in Na^+^ channel density of the lateral membrane on epicardial action potential parameters

## Results

### Reducing Na^+^ channel conductance in the LM causes AP morphological changes

First, we examined effects of alterations in subcellular Na^+^ channel expression on AP propagation by using a homogeneous myocardial strand model consisting of 300 cells. When the Na^+^ channel conductance of the LM (%*g*_Na,LM_) in each myocyte of the myocardial fiber was set to the control condition, the APs propagated through the myocardial strand (Fig. 2A*a*) and exhibited a typical human ventricular AP morphology (Figs. 2A*a* and 2B*a*, red line). To test how the selective decreasing of Na^+^ channel expression on the LM of myocytes as demonstrated by Shy et al (*10*) affects AP propagation, the %*g*_Na,LM_ in each myocyte of the myocardial fiber was reduced homogeneously. Reducing the %g_Na,LM_ by 65% resulted in the decrease in the *I*_Na,LM_ peak in the 150th cell by 57.9% (Fig. 2B*b*, cyan line) and slightly shortened the APD; APD at 90% repolarization (APD_90_) was 235.2 ms in 35%*g*_Na,LM_ versus 240.0 ms in 100%*g*_Na,LM_ (Figs. 2A*b* and 2B*a*, and Tables 1 and 2). These results were similar to those experimentally determined by the previous study (*8*, see table S3). Markedly reducing Na^+^ channels from the LM of each myocyte (7%*g*_Na,LM_) resulted in the *I*_Na,LM_ decrease by 92.5% (Fig. 2B*b*, blue line), diminished AP phase-0 amplitude and produced a larger phase-1 dip in the AP. As a result, the peak of the phase-2 dome was markedly delayed (Figs. 2A*c* and 2B*a*, blue lines). APD_90_ that delayed the phase-2 dome peak was 15.8 ms longer than that of the control condition (100%*g*_Na,LM_, see Table 2). Moreover, the loss of Na^+^ channels in the LM of each myocyte (0%*g*_Na,LM_) caused a further decrease in the AP phase-0 amplitude (Fig. 2B*a*, green line), resulting in the failure of the transient outward K^+^ channel current (*I*_to_) and L-type Ca^2+^ channel current (*I*_CaL_) to activate normally (Figs. 2B*c* and 2B*d*, green lines). The transient activation of the rapid component of delayed rectifier K^+^ channel current (*I*_Kr_) and the Na^+^-K^+^ pump current (*I*_NaK_), and subsequent activation of the inward rectifier K^+^ channel current (*I*_K1_), the late peak of which appeared earlier, allowed for earlier repolarization (Figs. 2B*e*, 2B*g* and 2B*h*). The slow component of delayed rectifier K^+^ channel current (*I*_Ks_) hardly contributed to the repolarization (Fig. 2B*f*, green line). This, in turn, led to AP propagation with a shortened APD (Fig. 2A*d*).

**Fig. 2.**
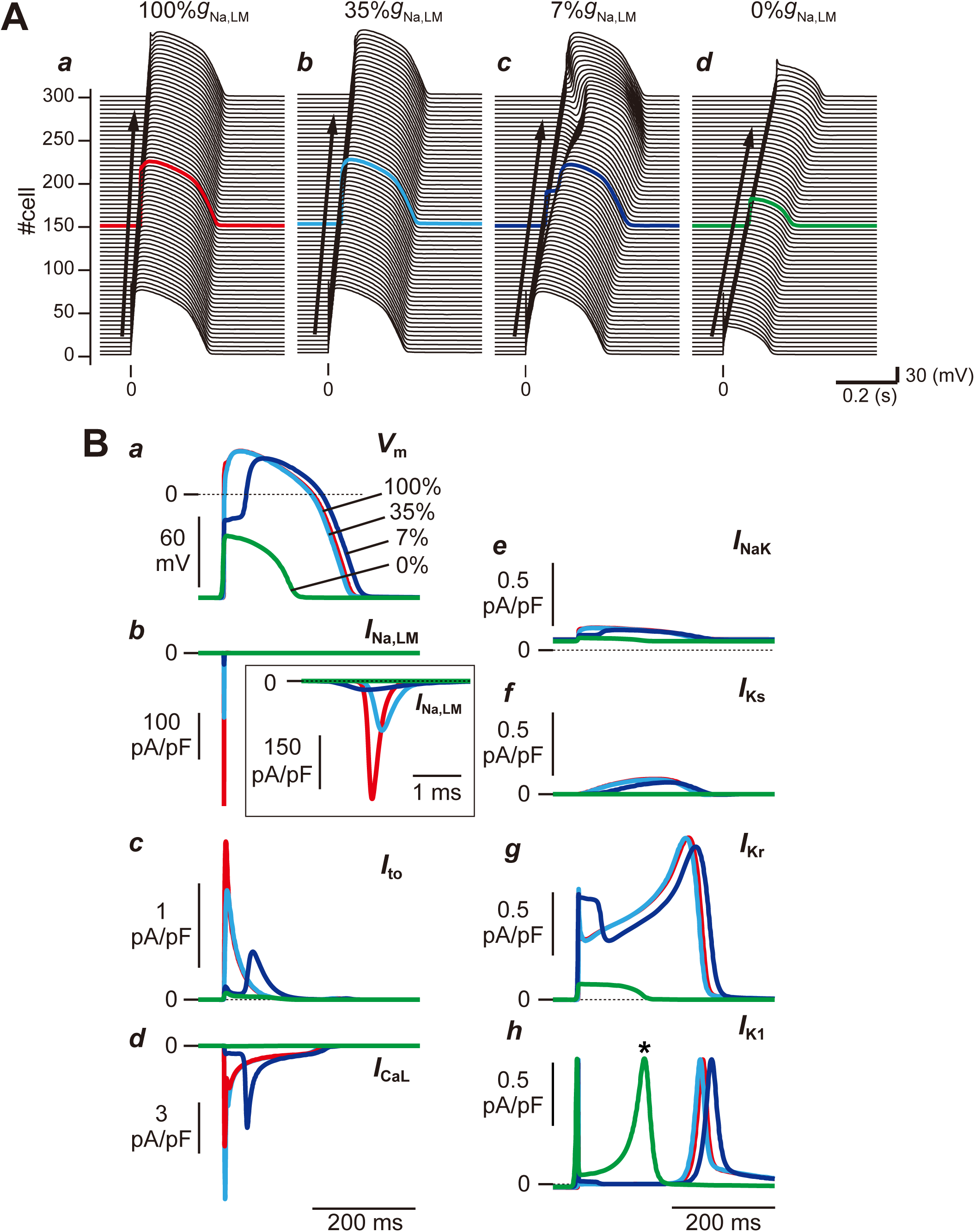
Effects of the subcellular Na^+^ channel distribution on action potential (AP) morphology and conduction velocity (CV). **A**, AP propagation observed in the myocardial strand model with a spatially-homogeneous reduction in Na^+^ channel conductance in the lateral membrane (LM) of 100%*g*_Na,LM_ (*a*), 35%*g*_Na,LM_ (*b*), 7%*g*_Na,LM_ (*c*), and 0%*g*_Na,LM_ (*d*). **B**, AP morphological changes (*a*), *I*_Na_ at the LM, *I*_Na,LM_ (*b*), *I*_to_ (*c*), *I*_CaL_ (*d*), *I*_NaK_ (*e*), *I*_Ks_ (*f*), *I*_Kr_ (*g*), and *I*_K1_ (*h*) at the LM. The APs and ion currents were recorded in a myocyte located at the middle of the myocardial strand (cell #150). Each AP morphology corresponds to the waveforms in panel (A) indicated by red, cyan, blue, and green lines. CVs at 100%*g*_Na,LM_, 35%*g*_Na,LM_, 7%*g*_Na,LM_ and 0%*g*_Na,LM_ were 71.4, 53.6, 33.3, and 25.4 cm/s, respectively (see Table 1).

These alterations in subcellular Na^+^ channel distribution also modified the conduction velocity (CV). The pre-JM current (*I*_m,pre-JM_; figs. S2A and S2B, orange lines) is the difference between the post-JM current (*I*_m,post-JM_; fig. S2B, purple lines) and the transmembrane current in the LM (see fig. S2C, gray lines), and is determined by *G*_i_ and *V*_i,LM_ − *V*_i,pre-JM_, where *V*_i,LM_ and *V*_i,pre-JM_ are the intracellular potentials in the LM and pre-JM compartments, respectively (see figs. S2A and S2D). The decrease in *I*_Na,LM_ (%*g*_Na,LM_ from 100% to 0%) caused a decrease in the maximum upstroke velocity of *V*_i,LM_ (compare gray lines in left to right panels of fig. S2D and see Table 2). The decrease in *V*_i,LM_ − *V*_i,pre-JM_ also reduced *I*_m,pre-JM_ (compare orange lines in left to right panels of fig. S2B), resulting in the decrease in maximum upstroke velocity of *V*_i,pre-JM_, ^msx^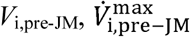, (compare orange lines in left to right panels of fig. S2D). Thus, the, 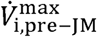 decrease in J _,bL6-}W_ caused a decrease in the difference between the *V*_i,pre-JM_ and the intracellular potential of the post-JM compartment (*V*_i,post-JM_; purple lines in fig. S2D) in the neighbor myocyte. Subsequently, this led to both a marked decrease in the gap junctional current (*I*_g_) defined by the equation 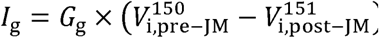 for 150th myocyte (compare colored lines in left to right panels of fig. S2E) and a decrease in CV (see Table 1).

### Combination of the decreases in Na^+^ channel expression on the LM in myocytes and the spatially-heterogeneous Na^+^ channel distribution in tissue led to P2R development

To test our hypothesis that alterations in the subcellular Na^+^ channel distribution lead to the development of P2R, we investigated the combined effects of both the regional *I*_Na_ decrease on the LM in myocytes and the spatial heterogeneity of Na^+^ channels in the myocardial tissue on AP propagation. The spatial heterogeneity of Na^+^ channels in the myocardial strand was achieved by changing the %*g*_Na,LM_ in the proximal part of the myocardial strand (200 myocytes), while maintaining the 7%*g*_Na,LM_ (0.77 nS/pF) in the distal part of the myocardial strand (100 myocytes). The AP conduction exhibited AP alternans (Fig. 3A*a*) when the %*g*_Na,LM_ of the proximal myocytes was set to 0.66 nS/pF (6%*g*_Na,LM_), as the slight spatial heterogeneity of Na^+^ channels in the myocardial strand. The spatial heterogeneity augmentation by further reducing the %*g*_Na,LM_ in the proximal myocytes led to proarrhythmic changes such that the AP dome was lost in the proximal myocytes and P2R followed (Fig. 3A*b*).

**Fig. 3.**
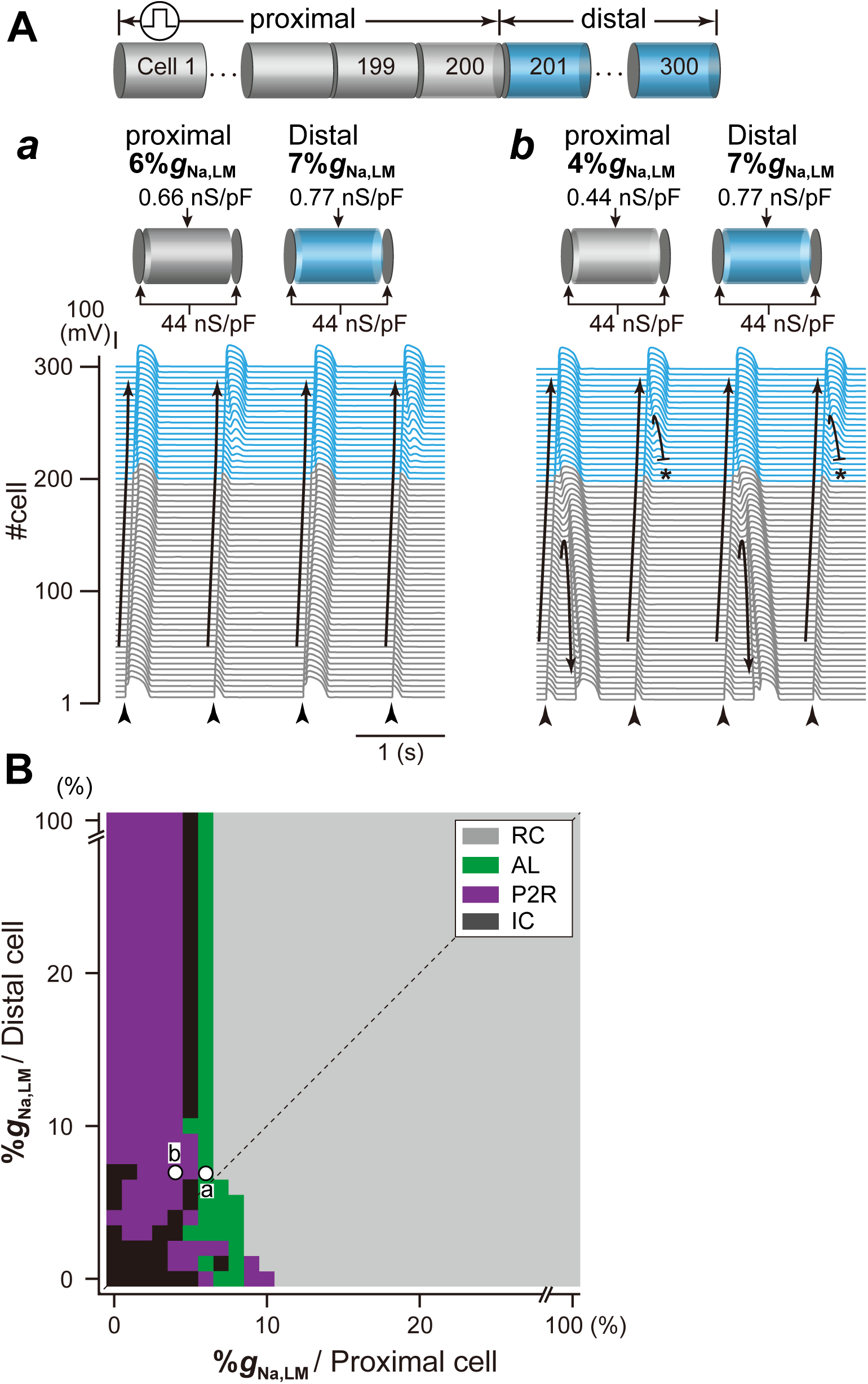
Simulated phase-2 reentry (P2R) development in the strand model. **A**, Examples of typical patterns of AP propagation observed in the myocardial strand model with a spatially-heterogeneous Na^+^ channel distribution on the LM of 7%*g*_Na,LM_ in the distal (cell #201-300) and smaller %*g*_Na,LM_ in the proximal (cell #1-200): AP alternans (AL) (*a*) and 2:1 retrograde P2R (*b*) at 6% and 4%*g*_Na,LM_, respectively, in the proximal. Arrows and black short bars indicate the direction of AP propagation and blockage, respectively. **B**, A phase diagram of AP propagation patterns for the %*g*_Na,LM_ in proximal myocytes vs. %*g*_Na,LM_ in distal myocytes. Open circles labeled as *a* and *b* correspond to the parameter sets for A(*a*) and A(*b*). RC and IC indicate AP propagation with regular pattern and irregular one, respectively.

To understand relationships among Na^+^ channel distributions within myocytes, spatial heterogeneous Na^+^ channel distribution in tissue, and development of P2R, we systematically performed simulations of AP propagations while changing the subcellular Na^+^ channel distribution in the proximal and distal parts of the myocardial fiber every 1%*g*_Na,LM_. Fig. 3B shows a phase diagram constructed by mapping the AP propagation patterns obtained from each simulation on the parameter plane of the %*g*_Na,LM_ in proximal and distal myocytes. Open circles labeled as “*a*” and “*b*” in Fig. 3B correspond to Figs. 3A*a* and 3A*b*, respectively. We can see that the 2:1 retrograde P2R as shown in Fig. 3A*b* occurred when the %*g*_Na,LM_ in the proximal myocyte was reduced to < 5% regardless of the %*g*_Na,LM_ in the distal myocytes (Fig. 3B, purple region), and that the region that P2R occurs was largely the left upper region of the diagonal line which represents conditions of spatially-homogeneous Na^+^ channel distribution in myocardial strand. This implies that P2R occurs when a spatially-heterogeneous Na^+^ channel distribution exists on the myocardial fiber. These results suggest that selective decrease in Na^+^ channels on the LM of myocytes and the spatially-heterogeneous Na^+^ channel expression on the myocardial tissue cooperatively facilitate the occurrence of P2R.

### I_CaL_ plays an important role in the P2R development

Based on the results shown in Fig. 3, we focused on how AP morphology is affected by changes in the ion channel currents. Fig. 4 shows the AP profiles and main ion channel currents in several myocytes near the border between the proximal and distal parts of the myocardial fiber during the development of the 2:1 P2R shown in Fig. 3A*b*. The initial depolarization of the regional membrane potential (*V*_m_) in the LM segment of each myocyte, i.e., that is equal to *V*_i,LM_, within the proximal part of the myocardial strand (#140, 160 and 180 in Fig. 4B, asterisk) was formed by the local current flows consisting of *I*_g_ (Fig. 4C) and post-*I*_Na,JM_ (Fig. 4D*a*). These current flows entered the myocyte through the post-JM. However, the local current was not able to depolarize the LM sufficiently (Fig. 4B, green line), and both *I*_CaL_ and *I*_to_ failed to activate (Fig. 4E*a-b*, green lines). Moreover, the relatively large transient *I*_Kr_ activation (Fig. 4E*c*, green line) and faster onset of the late peak of *I*_K1_ in the proximal myocyte (Fig. 4E*d*, dagger) accelerated the AP repolarization with the loss of AP dome (Figs. 4A and 4B).

**Fig. 4.**
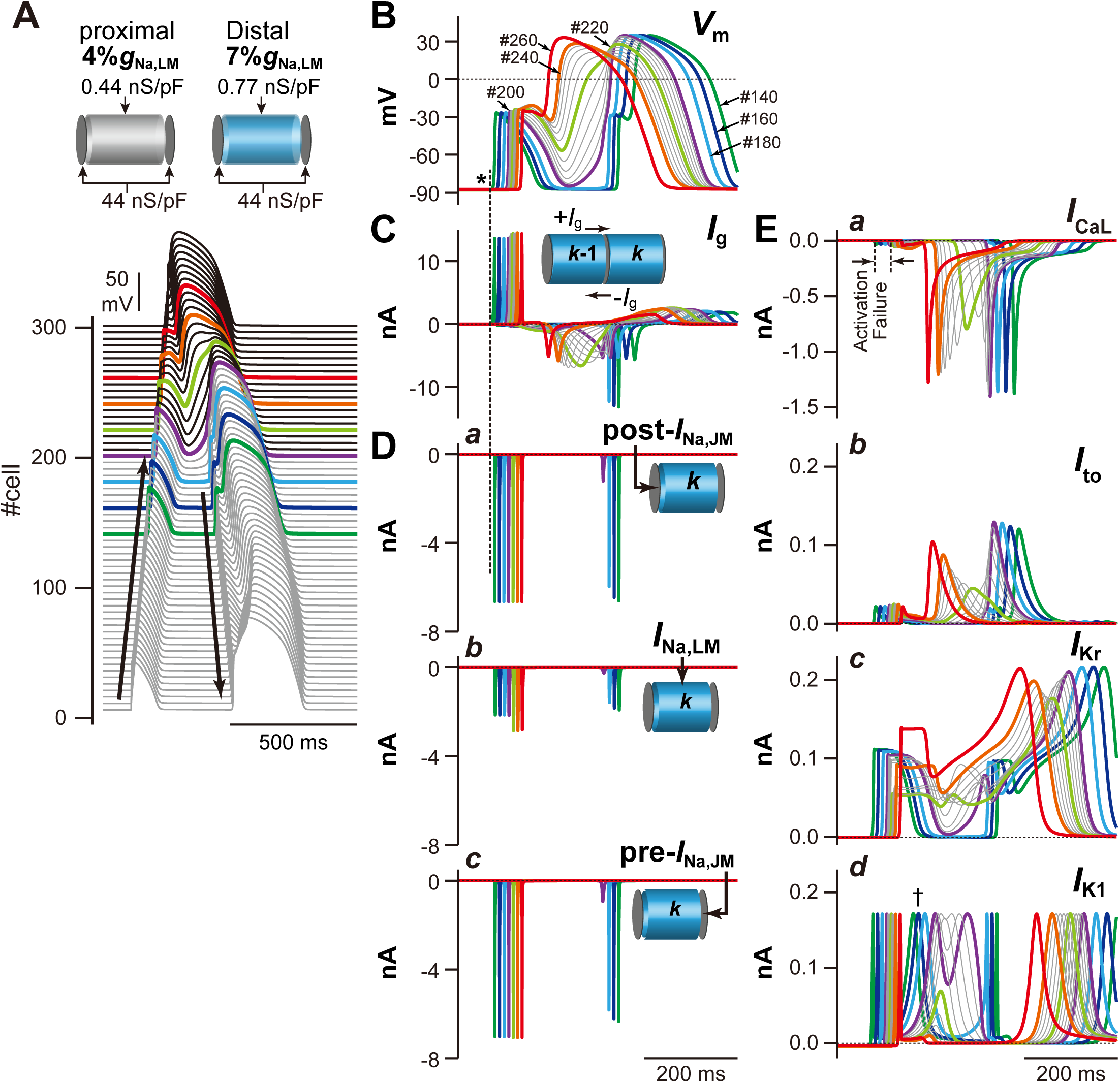
Mechanism of phase-2 reentry (P2R). **A**, The simulated AP propagation in the myocardial strand model during the development of 2:1 P2R shown in Fig. 3Ab. **B**, Simulated behaviors of the membrane potential (*V*_m_) in several myocytes (#140-260) during the 2:1 P2R development. **C**, The gap junctional currents, *I*_g_, flowing from the cell #140 to #260. **D**, *I*_Na_ in the post-junctional membrane (*I*_Na,post-JM_) (*a*), the lateral membrane (*I*_Na,LM_) (*b*), and the pre-junctional membrane (*I*_Na,pre-JM_) (*c*). **E**, *I*_CaL_ (*a*), *I*_to_ (*b*), *I*_Kr_ (*c*), and *I*_K1_ (*d*) during the P2R development.

Subsequently, the difference in *V*_m_ at the border between proximal and distal parts was increased, and consequently this elicited the retrograde *I*_g_ (the inward currents in Fig. 4C) from the distal to the proximal part. Unlike the anterograde *I*_g_ elicited by the initial depolarization differences in neighbor myocytes within the proximal part, the retrograde *I*_g_ continued to flow for a relatively long time because the notch-and-dome type prolonged AP in distal myocytes caused a long-lasting potential difference from the proximal loss-of-dome abbreviated AP. Continuous depolarizing loads of the proximal myocytes by the long-lasting retrograde *I*_g_ allow the depolarization of the *V*_m_ that activates *I*_CaL_ of the LM in proximal myocytes, leading to re-excitation of the myocytes in the proximal part, i.e., retrograde P2R (Fig. 4A).

This theoretical model for the P2R mechanism can also be used to investigate the mechanism of P2R inhibition. We conducted an additional, identical simulation under a *β*-AS condition (*20, 21*). The marked increase in *I*_CaL_ due to the *β*-AS effect kept the AP dome of each myocyte even under the smaller %*g*_Na,LM_ condition that caused loss-of-dome type abbreviated AP by the marked decrease in *I*_Na,LM_ in the proximal part of the myocardial fiber without *β*-AS (see Fig. 5). This resulted in an increase in *V*_m_ amplitude and APD prolongation in the proximal and distal parts of the myofiber (compare Fig. 5B and Fig. 4B). Thus, *β*-AS that causes an increase in *I*_CaL_ is effective for preventing P2R.

**Fig. 5.**
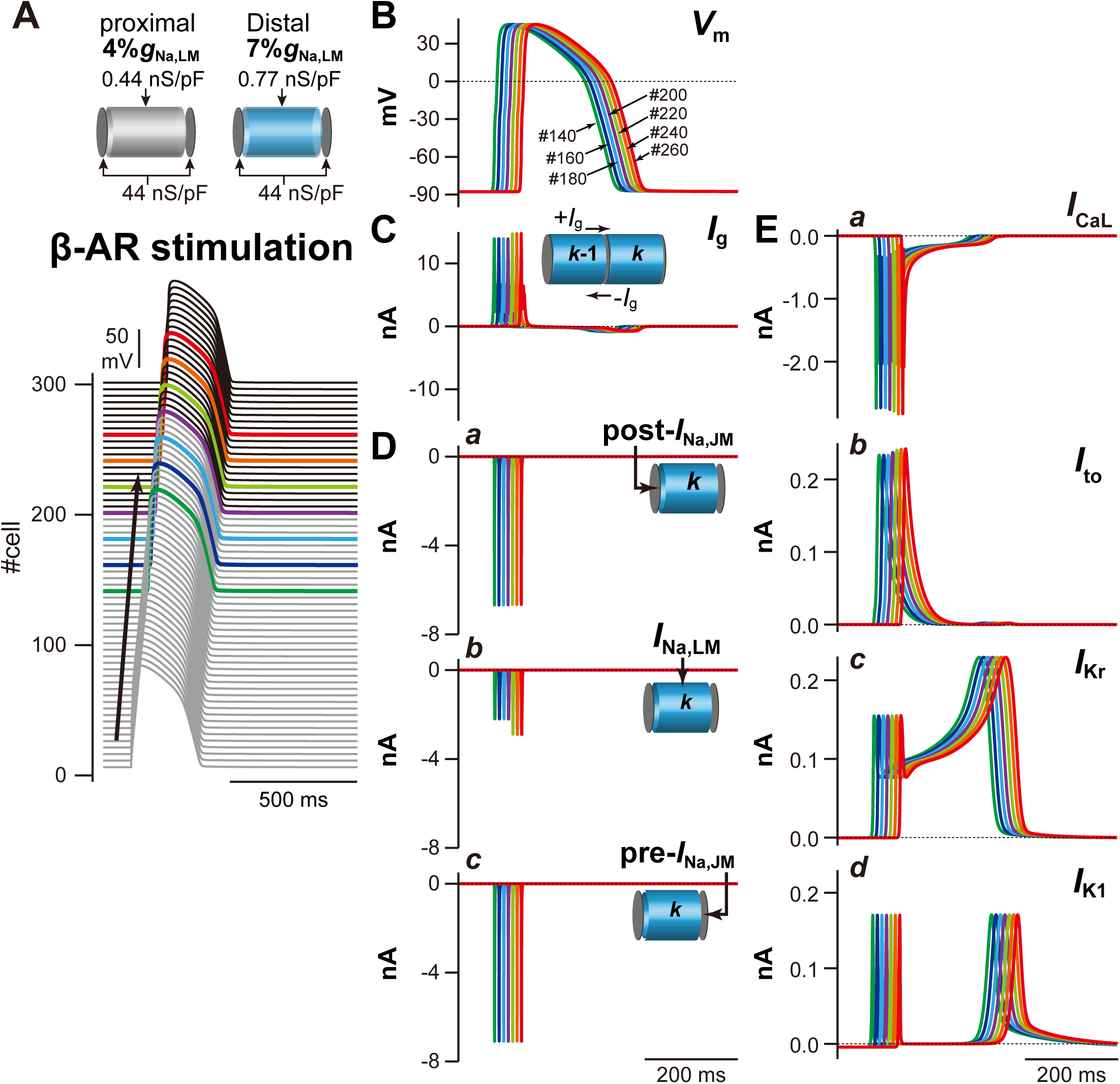
Mechanisms for the prevention of phase-2 reentry (P2R) under *β*-AS condition. **A** and **B**, The simulated AP propagation (A) and the membrane potential (*V*_m_) change (B) of several myocytes in the myocardial strand model under the same condition as for Fig. 3Ab but during *β*-AS. **C**-**E**, The gap junctional current, *I*_g_ (C), *I*_Na_ in the post-junctional membrane (JM), *I*_Na,post-JM_ (D*a*), *I*_Na_ in the lateral membrane, *I*_Na,LM_ (D*b*), and *I*_Na_ in the pre-JM, *I*_Na,pre-JM_ (D*c*), *I*_CaL_ (E*a*), *I*_to_ (E*b*), *I*_Kr_ (E*c*), and *I*_K1_ (E*d*) during *β*-AS.

### The P2R triggered reentrant arrhythmias

Whether or not the P2R triggers reentrant arrhythmias was investigated using a myocardial ring model. In the myocardial ring with spatially-homogeneous Na^+^ channel distribution (Fig. 6A), the AP exhibited bidirectional conduction from the stimulus site (300th myocyte), resulting in the collision of excitation wavefronts at the opposite side of the stimulus site (Fig. 6A*a*, asterisk). When the %*g*_Na,LM_ from the LM of each myocyte in the myocardial ring was homogeneously reduced by 95%, the bidirectional conduction of shortened AP from the stimulus site was observed (Fig. 6A*b*); the bidirectional excitation wavefronts collided, causing a bidirectional P2R in the myocardial ring. However, each excitation wavefront originating from the bidirectional P2R collided and disappeared at around stimulus site (Fig. 6A*b*); no proarrhythmic changes were observed in the myocardial ring model with spatially-homogeneous Na^+^ channel distribution even under the condition of the markedly decreased %*g*_Na,LM_. However, in the same myocardial ring model but with the spatially-heterogeneous Na^+^ channel distribution (Fig. 6B), AP conduction properties obviously changed (see Figs. 6B*a* and 6B*b*). With maintaining the %*g*_Na,LM_ of 100 cells (cells #1-100; region A in Fig. 6B) at 5% (0.55 nS/pF), the %*g*_Na,LM_ in each 500 cells (cells #101-600; region B in Fig. 6B) was uniformly reduced to 3% (0.33 nS/pF), producing the spatial heterogeneity of Na^+^ channel distribution within the tissue. The bidirectional excitation wavefronts elicited at the stimulus site (cell #300) collided at the border between the regions A and B (Fig. 6B*a*, asterisk) and were subsequently followed by bidirectional P2R occurrence. However, the clockwise-rotating excitation wave of the P2R exhibited decremental conduction and block (Fig. 6B*a*, dagger); in contrast, the counterclockwise-rotating excitation wave continued to conduct in the myocardial ring, causing the P2R-mediated reentrant tachyarrhythmias. During 1-Hz pacing, the occurrence of reentrant arrhythmias was intermittent and irregular (see fig. S3A). The further augmentation of the spatial heterogeneity of Na^+^ channel distribution in the tissue (5%*g*_Na,LM_ in region A and 2%*g*_Na,LM_ in region B on the myocardial ring) induced P2R-mediated persistent reentry (Figs. 6B*b* and S3B).

**Fig. 6.**
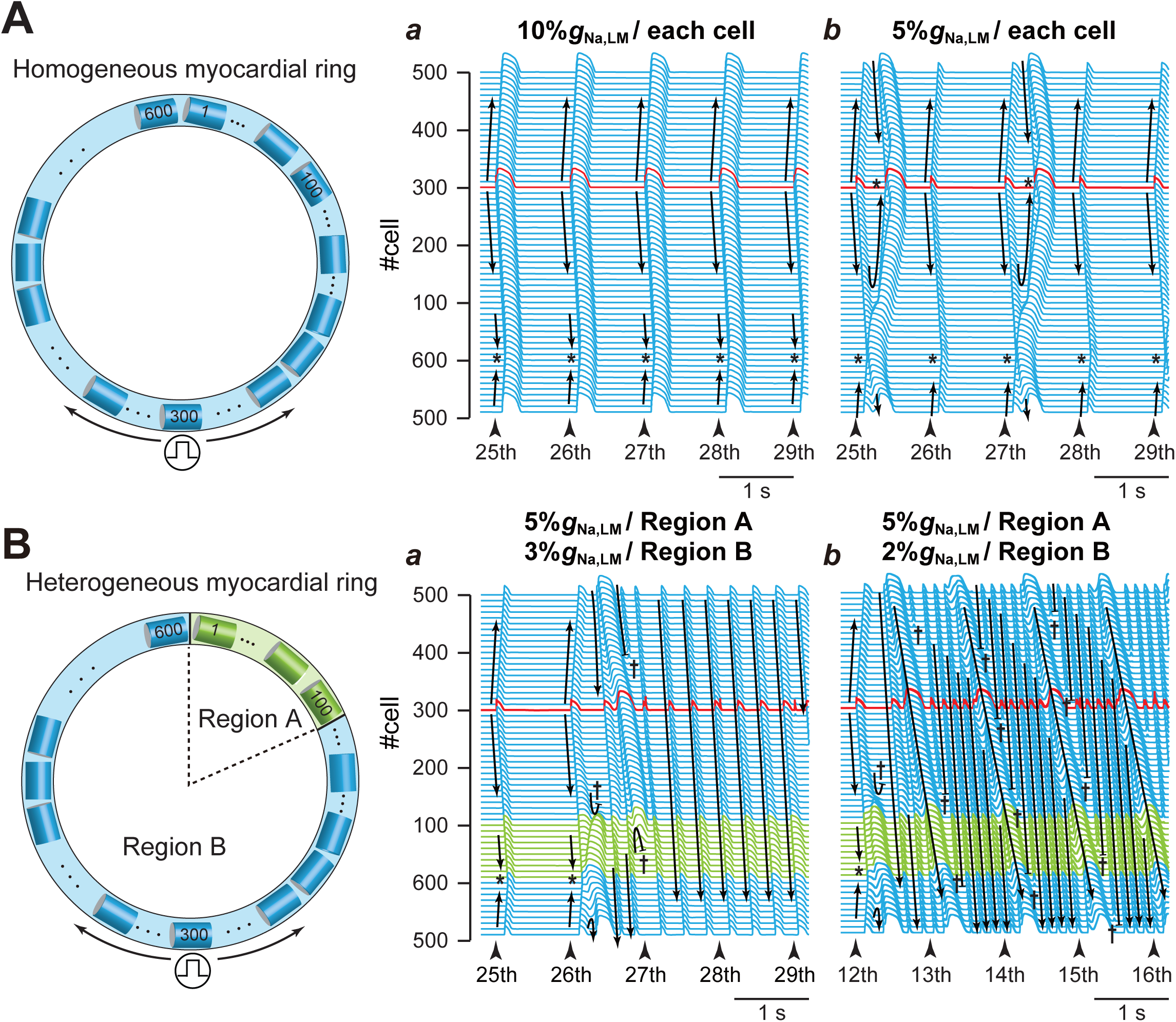
Rotatory reentry induction in the myocardial ring model. **A**, The simulated AP propagation in the myocardial ring with spatially-homogeneous Na^+^ channel distribution with 10%*g*_Na,LM_ (*a*) and 5%*g*_Na,LM_ (*b*) in the lateral membrane (LM) segment of each myocyte. **B**, The simulated AP propagation in the same model but with a spatially-heterogeneous Na^+^ channel distribution with 5%*g*_Na,LM_ in the LM of the 1st to 100th myocytes and 3%*g*_Na,LM_ (*a*) and 2%*g*_Na,LM_ (*b*) in the LM of the 101st to 600th myocytes. Arrows and black short bars indicate the direction of AP propagation and blockage, respectively. Asterisks and dagger symbols represent the collision of excitation wavefronts, and the blockade of AP propagation, respectively. The red trace represents the AP behavior at the stimulus site (cell #300).

## Discussion

Although it has so far believed that the loss-of-function mutation of Na^+^ channels is one of the major causes of fatal arrhythmias in patients with BrS, the link between the loss-of-function of Na^+^ channels and arrhythmogenesis was not clear. In the present study, we found that selective decreases in *I*_Na,LM_ of each ventricular myocyte caused both morphological changes in AP and the conduction slowing that are typically observed in BrS patients (*22, 23*). These findings suggest that both the BrS mechanisms, previously represented as repolarization and depolarization abnormalities (*24*), can at least in part be explained by this decrease in *I*_Na,LM_. Furthermore, we showed that the marked decrease in *I*_Na,LM_ together with an existence of the spatial heterogeneity at tissue-level in the Na^+^ channel expression is essential for the P2R development, as well as the reentry initiation triggered by the P2R. To the best of our knowledge, this is the first report demonstrating mechanistic links between subcellular Na^+^ channel expression changes mediated loss-of-function of Na^+^ channels and P2R in BrS.

### Decreasing I_Na,LM_ causes the BrS notch and dome AP morphology

An alteration in the subcellular Na^+^ channel expression with the marked reduction in Na^+^ channels in the LM (%*g*_Na,LM_ to 7%) resulted in the notch-and-dome AP morphology (Fig. 2A*c* and 2B*a*, blue line). Selectively reducing Na^+^ channels from the LM largely reduced the AP phase-0 amplitude (Fig. 2B*a*, blue line). Because both *I*_to_ and *I*_CaL_ activate as *V*_m_ exceeds −30 mV (*16*), the underdeveloped AP phase-0 amplitude resulted in a marked decrease in *I*_to_ during AP phase-1 and the delay of *I*_CaL_ activation (Figs. 2B*c* and 2B*d*, blue lines). The decrease in *I*_to_ and delay in *I*_CaL_ activation led to a large phase-1 dip and the delay of the AP phase-2 dome peak, respectively. This morphology resembled that of epicardial monophasic AP recordings from the RV outflow tract (RVOT) in patients with BrS who were undergoing open chest surgery (*22*). This implies that the characteristic monophasic AP morphology recorded in BrS might be attributed to the selective decrease of *I*_Na,LM_ in epicardial myocytes located in the RVOT.

### Loss of I_Na,LM_ causes the BrS loss-of-dome AP morphology

The loss of the AP dome and marked shortening of APD in epicardial myocytes in the RVOT, i.e., repolarization abnormality (*6, 24*), are considered to be one of the mechanisms of coved-type ST-segment elevation (Brugada-type ECG) (*25*). One of the factors causing the loss-of-dome is thought to be the ventricular gradient of *I*_to_ through the RV wall, as well as endogenously heterogeneous *I*_to_ in the RV epicardial layers (*6*). We showed that the loss-of-dome type abbreviated AP was also caused by extremely decrease and/or loss of Na^+^ channels from the LM of each myocyte, even without changing *I*_to_ (Figs. 2A*d* and 2B*a*; green lines). Indeed, the loss of *I*_Na,LM_ further decreased the AP phase-0 amplitude, therefore preventing the *V*_m_ depolarization at AP phase-0 from reaching −30 mV, a threshold potential for *I*_CaL_ activation (see Fig. 2B*a*, green line); this thus failed to activate *I*_CaL_ (Fig. 2B*d*), resulting in the loss-of-dome type abbreviated AP (Fig. 2B*a*, green line). Based on the present simulation study, the loss of the AP dome can be also explained by both the *I*_Kr_ activation (Fig. 2B*g*, green line) and the earlier late peak (*) of *I*_K1_ (Fig. 2B*h*, green line), even without the activation of *I*_to_ (please note that the very small *I*_to_ was observed in Fig. 2B*c*). These results suggest that the Na^+^ channel abnormality, which is commonly believed to be the cause of the depolarization abnormality in BrS (*24*), may also be responsible for the repolarization abnormality of BrS.

### Selective decrease in I_Na,LM_ causes the depolarization abnormality in BrS

We found that the conduction delay in RVOT under structurally normal hearts in patients with BrS (*26*), i.e., depolarization disorder hypothesis (*24*), may be attributable to the selective decrease in *I*_Na,LM_ causing decreases in *I*_g_ without the *G*_g_ change. The mechanism of *I*_Na,LM_-mediated decreases in *I*_g_ proposed in the present theoretical study is as follows: (1) The marked decrease in *I*_Na,LM_ decreases the pre-JM current (compare left to right panels of fig. S2C) and thus slows the depolarization of the pre-JM (compare left to right panels of fig. S2D). (2) The slower and smaller depolarization of the pre-JM due to the reduced pre-JM current leads to decreases in the difference between 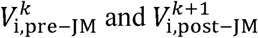, for *k* = 1, 2,…, 299, resulting in the *I*_g_ decrease. This notion is also supported by previous experimental studies (*9, 10*), which showed that mutant mice hearts with a selective decrease in Na^+^ channel expression at the LM had slower CV than the wild type, without affecting the connexin 43 (Cx43; gap-junction channel-forming proteins) expression (table S3).

### Possible mechanisms of P2R and related arrhythmias

The underlying mechanism of P2R is commonly explained by the electrotonic interaction between the region with normal AP morphology and the adjacent region with the loss of the AP dome. Such P2R mechanism were explored in numerous experimental (*27, 28*) and computational studies (*29–31*). For the onset of P2R, excitable tissues such as RV and RVOT, in general, are required to be separated into two regions by the segment of reduced excitability and/or reduced electrical coupling. The existence of a local difference in AP waveform (or local APD difference) at the RVOT epicardium has been confirmed by several experimental studies (*8, 32*). Our results showed that an existence of the slight spatial heterogeneity at tissue-level in Na^+^ channel expressions accompanied by a marked decrease in *I*_Na,LM_ (reduced excitability) could induce P2R (Fig. 3). Although the causes that make the local differences of APs in the RVOT/RV are not clear, one possible cause is an endogenous heterogeneity in Na^+^ channel expression at the RVOT epicardial layer as in the transmural difference in Na^+^ channel expression in the RVOT and the RV free wall (*33*). On the other hand, Auerbach and co-workers (*34*) have shown by combined experimental and computational studies that a structural heterogeneity (a geometric expansion) can produce P2R. In addition to differences in electrophysiological properties due to differences in the developmental origin of the RVOT and RV (*35*), a tissue structure of the RVOT with more aligned fiber orientations is different from that of the RV free wall which comprised of network-like structures of myocardial fibers; the structural heterogeneity between the RV free wall and RVOT might be also responsible for the P2R development in patients with BrS.

A recent clinical observation (*36*) has demonstrated that BrS is associated with epicardial interstitial fibrosis and reduced Cx43 expression in the RVOT. Furthermore, previous experimental studies using the *Scn5a* heterozygous (*Scn5a+*/-) mice (*37, 38*) have shown that fibrosis, accompanied by downregulation and redistribution of Cx43, is increased with aging. The *I*_g_ decrease with *G*_g_ change leads to a decrease in the local current (*I*_m,LM_) consisting of *I*_g_ and *I*_m,post-JM_ (figs. S2B and S2E). Besides *I*_Na,LM_ reduction, decreases in *I*_g_ could result in the loss-of-dome abbreviated AP (*31*). Therefore, we speculate that the decrease in gap junctions at the RVOT, where Na^+^ channel expression is impaired along with aging-associated fibrosis, may also cause the loss of the AP dome followed by P2R resulting in VT/VF.

### Contribution of I_CaL_ to P2R

Miyoshi et al (*30*) suggested that *I*_CaL_ plays critical roles in the formation of local APD differences even during AP propagation and the following second excitation during P2R. Indeed, as shown in Figs. 2B and 4B, *I*_CaL_ was the main determinant of AP dome formation in each myocyte of the myocardial strand. As the original ORd model was constrained as AP behaviors observed in the human left ventricular endocardial myocyte, actual *I*_CaL_ in the RVOT might be different from that in the left ventricular myocyte. In the present study, we increased the *I*_CaL_ by 30% from the original value in the ORd model (*16*) to reproduce P2R development. This modification was needed to maintain the second excitation wave propagation during P2R, consequently leading to reentrant excitation wave propagations (VT/VF) in a myocardial ring model (Fig. 6).

On the other hand, it is known that the sympathetic activation in patients with BrS decreases VT/VF initiation (*39*), predicting that the greater *I*_CaL_ enhancement contributes to preventing P2R. We demonstrated that *β*-AS by the administration of isoproterenol could reduce the local APD difference and could prevent P2R (Fig. 5); more quantitative experimental verification and theoretical studies using more elaborate intracellular Ca^2+^-signaling models (*40*) are required to understand the exact roles of *I*_CaL_ modulation by *β*-AS in P2R and arrhythmia preventions. Furthermore, additional simulations employing more sophisticated, three-dimensional, whole ventricle models will be needed to elucidate the roles of the loss-of-function of Na^+^ channels in BrS.

## Author contributions

K.T., T.A. and Yo.K. conceived and designed the experiments; K.T., N.N., and T.S. conducted the simulations, performed numerical calculations and prepared figures; K.T., T.A. N.N., T.S., Ya.K., and Yo.K. analyzed and interpreted results; K.T., T.A., Ya.K., A.A. and Yo.K. drafted and edited the manuscript. All authors reviewed and approved the final version of manuscript. Competing interests: The authors declare no competing financial interests.

## Data and materials availability

These simulation codes of the myocardial strand and ring models are available on the author’s GitHub site: https://github.com/92tsumoto/BrS-P2R-strand-ORd2011model-withTNNP_INa-FT_IKr, and https://github.com/92tsumoto/BrS-P2R-ring-ORd2011model-withTNNP_INa-FT_IKr.

## Supplementary Materials

### Expanded Methods

#### Myocardial strand and ring models

We constructed myocardial strand and ring models comprising 300 (Fig. 1A) and 600 ventricular myocytes (Fig. 1D), respectively. As with our previous study (*11*), the cytosolic conductance (*G*_i_) of each myocyte was 2.327 μS, calculated from *G*_i_ = σ_myo_·π·*r*^2^/*l*, where σ_myo_ (11.1 mS/cm) is the cytosolic conductivity, and *l* (150 μm) and *r* (10 μm) are the length and radius of the myocyte, respectively (*12*). In addition, the radial cleft conductance (*G*_j_) and series axial cleft conductance (*G*_d_) were defined as *G*_j_ ·π· *r*^2^/*w*, respectively (*13, 14*), where σ_ext_ (6.7 mS/cm) is the extracellular conductivity, and *w* (15 nm) is the cleft width, which is the distance between the pre- (pre-JM) and post-junctional membrane (post-JM), as an averaged value (*11, 13*).

Electrophysiological properties of each membrane segment composing the ventricular myocyte were described by a modified O’Hara–Rudy dynamic (mORd) model (*16*). The mORd model consisted of a membrane capacitance and several ion channel currents, including our modification (*17*) to the original currents of a potassium channel (*I*_Kr_). Furthermore, the fast *I*_Na_ in the original ORd model was replaced with that of the ten Tusscher-Panfilov (TP) model (*19*). From experimental data (*12*), the entire membrane capacitance of a human ventricular myocyte was set to 185 pF (*C*_m,post-JM_ = *C*_m,pre-JM_ = 5.8 pF; *C*_m,LM_ = 173.4 pF) and the initial value of [Na^+^]_i_ was set to 7.0 mM. Table S1 shows the modification parameters of the relevant models.

### Calculations

The calculation methods for action potential propagation in the myocardial strand and ring models are as follows. From the equivalent circuits of the myocyte model shown in Fig. S1, let us give the following current vector (**I**) comprised of the transmembrane current from each node toward each membrane segment:

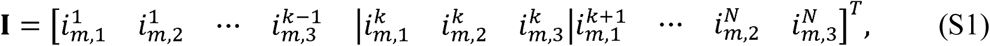

where []*^T^* represents the transpose operation and *N* is the total myocyte number. Furthermore, we suppose that a vector (**V**) consisting of the functions of the transmembrane potential of each membrane segment is given by

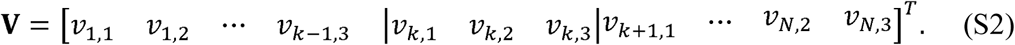

Then, each function, *v_k,l_*, for *k* = 1, …, *N* and *l* = 1, 2, and 3, except for the first function (*v*_1,1_) and the final one (*v_N_*_,3_), in the voltage vector is given by

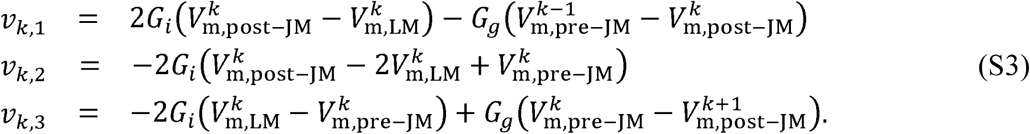

In cases of *k* = 1 and *k* = *N*, *v*_1,1_ and *v_N_*_,3_ are provided by 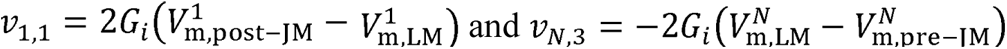, respectively. Using the current vector (Eq. S1) and the vector (Eq. S2) we can derive a linear equation at arbitrary time *t* using Ohm’s law and Kirchhoff’s law, leading to the following simultaneous equation:

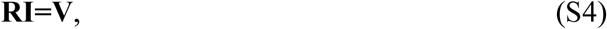

where **R** is the 3*N* × 3*N* matrix. The matrix, **R**, in Eq. S4 is given by Eq. S5:

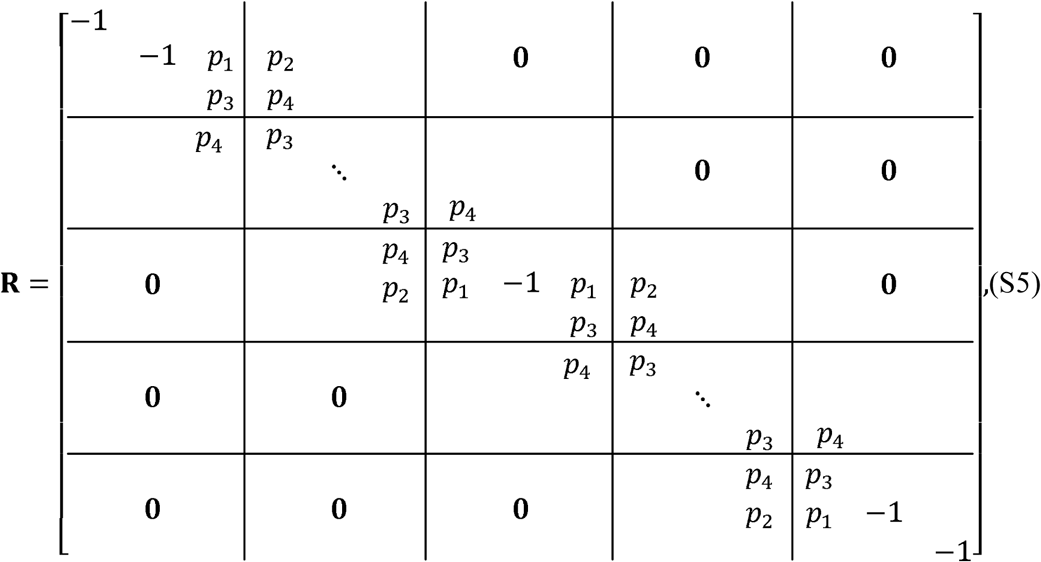

where *p*_1_ = 2*G*_i_/*G*_j_ + *G*_i_/*G*_d_, *p*_2_ = 2*G*_i_/*G*_j_, *p*_3_ = −1.0 − 2*G*_i_/*G*_j_ − *G*_i_/*G*_d_ − 0.5 *G*_g_/*G*_d_, and *p*_4_ = −2*G*_i_/*G*_j_ + 0.5 *G*_g_/*G*_d_. Equation S4 can be solved for the current ***I*** with 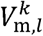, for *l* = post-JM, LM, and pre-JM, at time *t* as an initial condition. Furthermore, in the case of the myocardial ring model, the following matrix was employed:

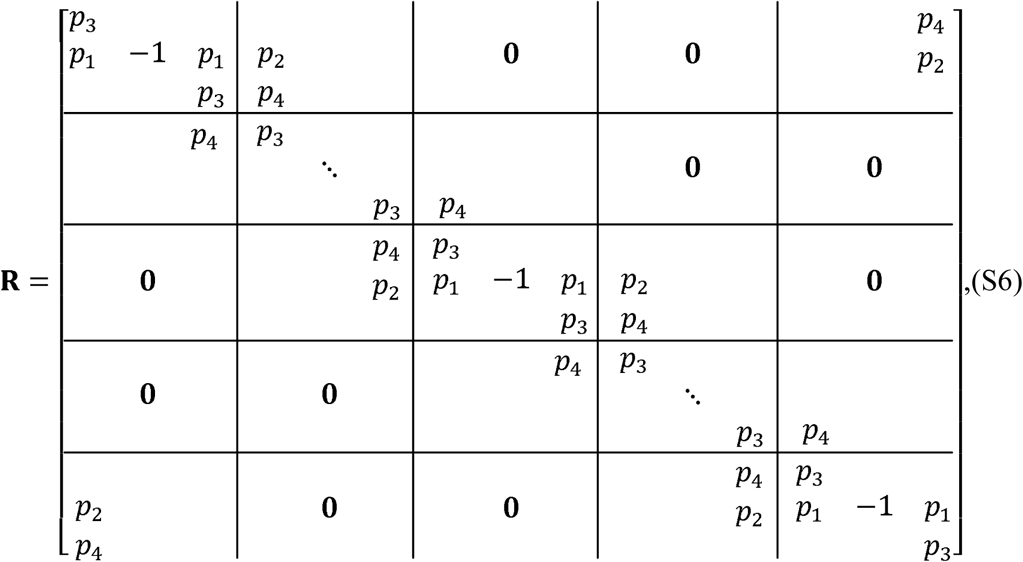

The transmembrane potential in each segment is given by

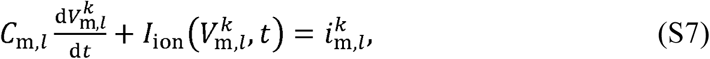

for *k* = 1, …, *N* and *l* = post-JM, LM, and pre-JM, where *C*_m,*l*_ (pF) is the membrane capacitance in each membrane segment, *I*_ion_ (μA/μF) is the sum of several ion channel currents in the mORd model, 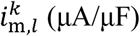 is the transmembrane current corresponding to each element in the current vector defined by Eq. S1. For an arbitrary time *t*, all the membrane currents, 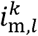, are obtained by solving Eq. S4 with the transmembrane potentials as an initial condition. Thus, we calculated all transmembrane potentials at time *t*+ Δ *t* in each segment, where Δ*t* corresponds to the s. These codes used to time step in the Euler method. The time step, Δ*t*, was set to 1 μ simulate the myocardial strand and ring models are available in the repository: https://github.com/92tsumoto/BrS-P2R-strand-ORd2011model-withTNNP_INa-FT_IKr, and https://github.com/92tsumoto/BrS-P2R-ring-ORd2011model-withTNNP_INa-FT_IKr.

## Supplemental Figures

**Fig. S1.**
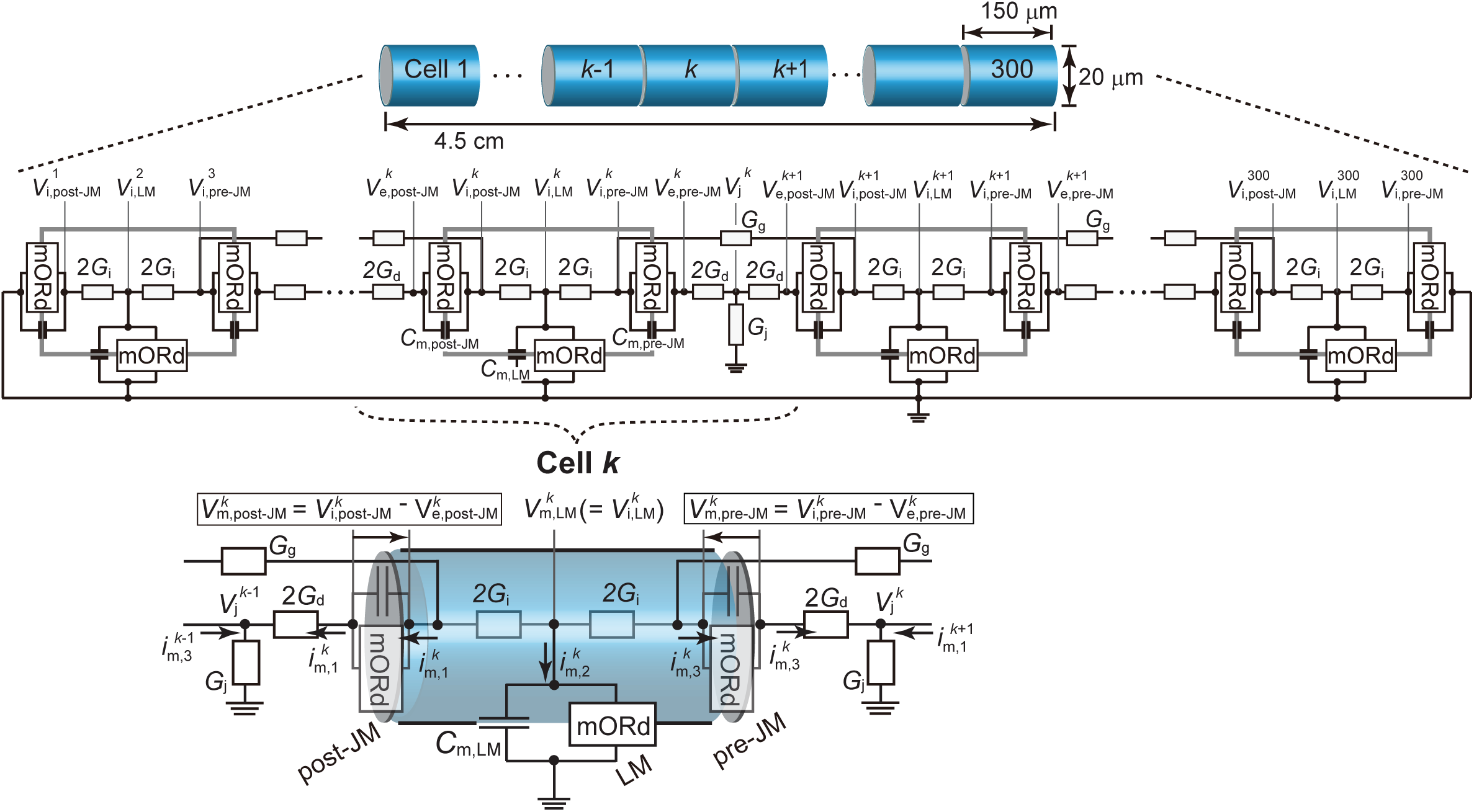
Myocardial strand model and its equivalent circuit. Each membrane segment comprises a modified O’Hara-Rudy dynamic (mORd) model and membrane capacitance, *C*_m,*l*_, for *l* = post-junctional membrane (post-JM), lateral membrane (LM), and pre-junctional membrane (pre-JM).*V_j_^k^* represents extracellular cleft potential just after the *k*th myocyte. 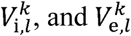, for *l* = post-JM, LM, and pre-JM, indicate the intracellular and extracellular potentials, respectively, of *l*th segment of the *k*th myocyte. denotes the transmembrane potential, i.e.,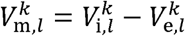

**Fig. S2.**
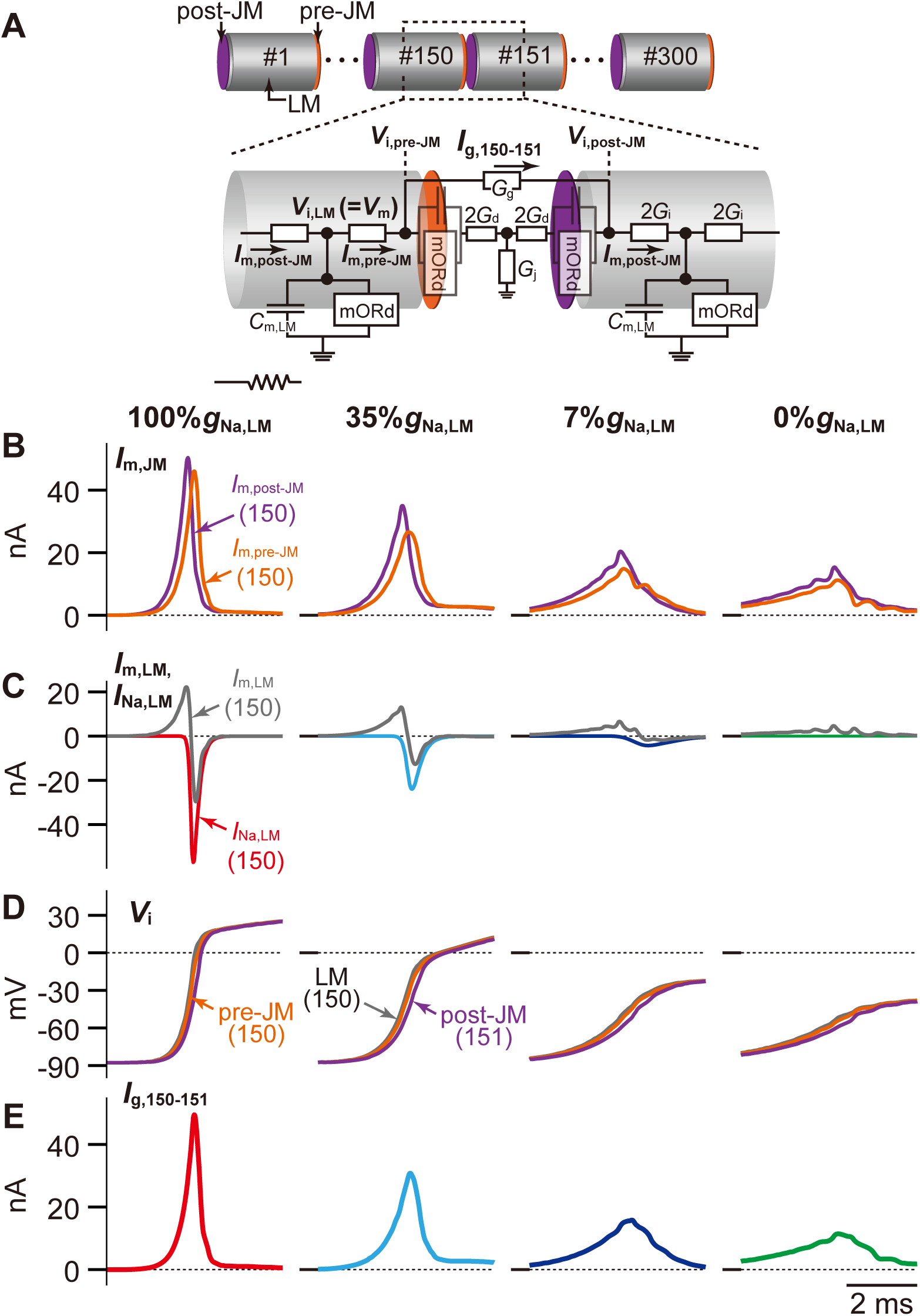
Effect of *I*_Na_ changes in the lateral membrane (LM) on conduction velocity (CV). **A**, The schematic diagram of a intercellular junction part in the myocardial strand model. **B**, The post-junctional (*I*_m,post-JM_) and pre-junctional transmembrane currents (*I*_m,pre-JM_) in 150th myocyte. **C**, The transmembrane current (*I*_m,LM_) and *I*_Na_ in the LM segment of 150th myocytes. **D**, The intracellular potential (*V*_i_) of the LM and pre-junctional membrane (pre-JM) in 150th myocyte, and of the post-junctional membrane (post-JM) in 151st myocyte. **E**, The gap junctional current (*I*_g_) between the 150th and 151st myocytes.

**Fig. S3.**
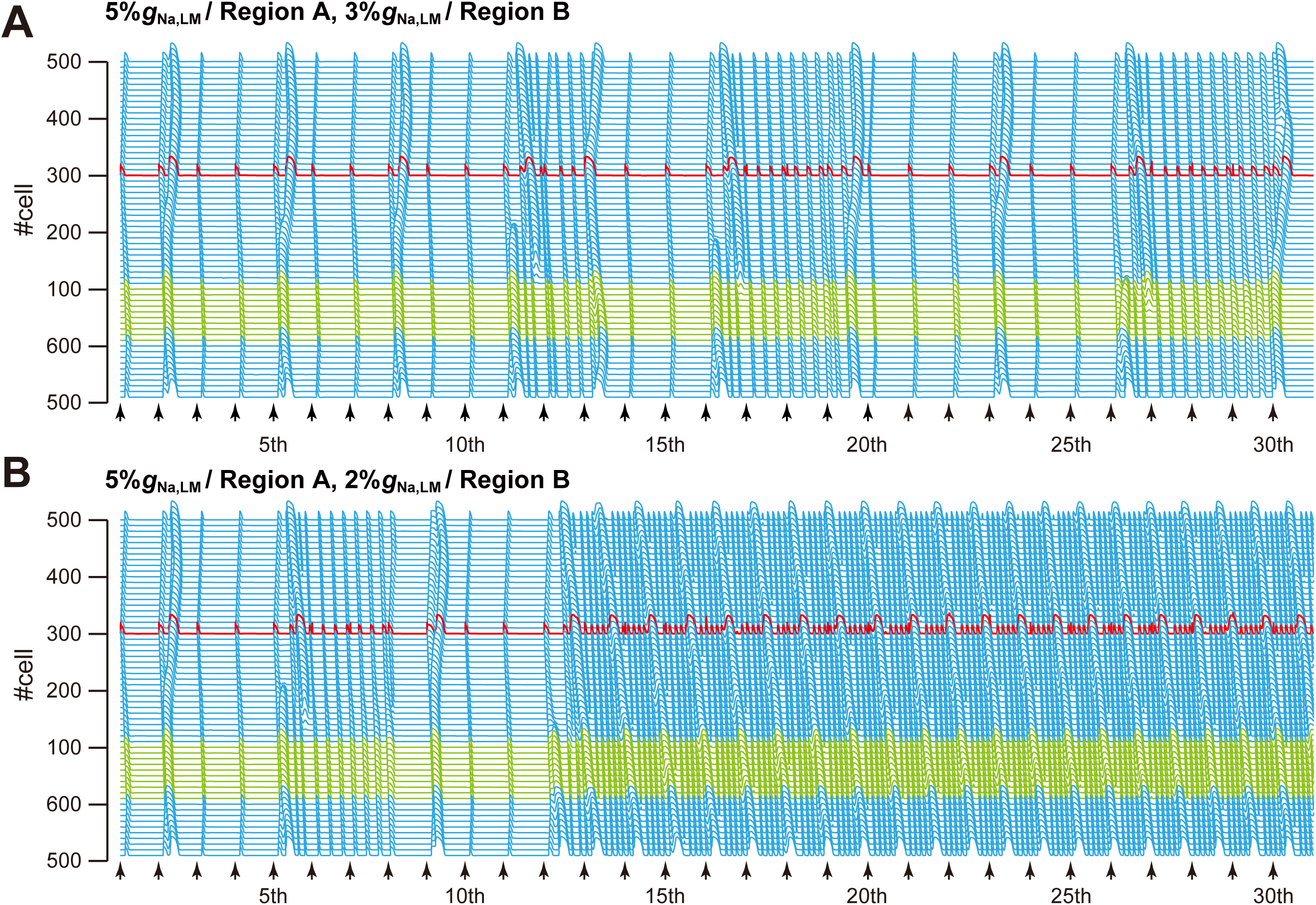
Overview of simulated AP propagation in myocardial ring models with spatially-heterogeneous Na^+^ channel distributions. **A**, 5%*g*_Na,LM_ in the LM of 1st to 100th myocytes and 3%*g*_Na,LM_ in the LM of 101st to 600th myocyte. **B**, 5%*g*_Na,LM_ in the LM of 1st to 100th myocytes and 2%*g*_Na,LM_ in the LM of 101st to 600th myocyte. Representations are the same as in Fig. 6.

## Supplemental Tables

**Table S1.**
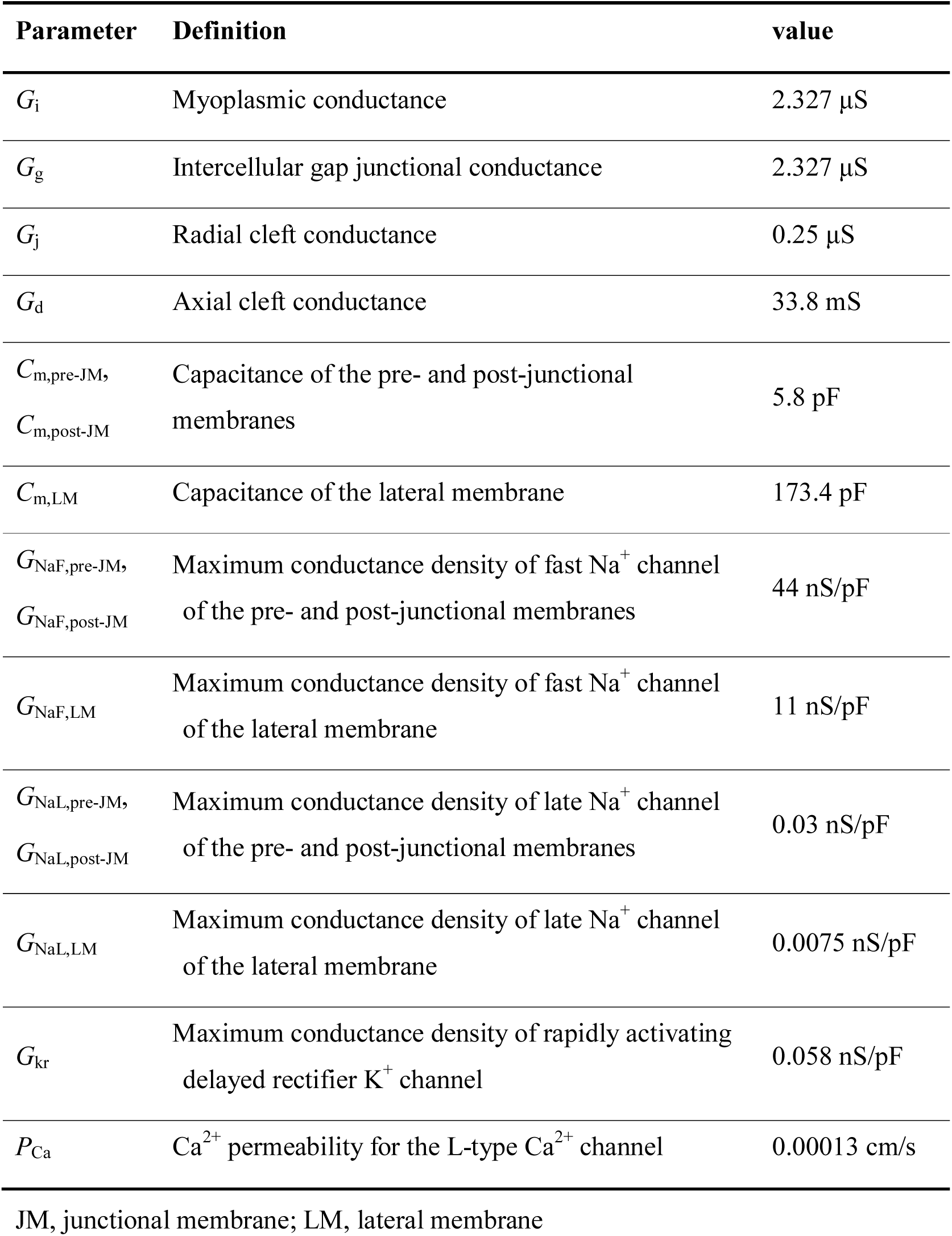
Control parameter values

**Table S2.**
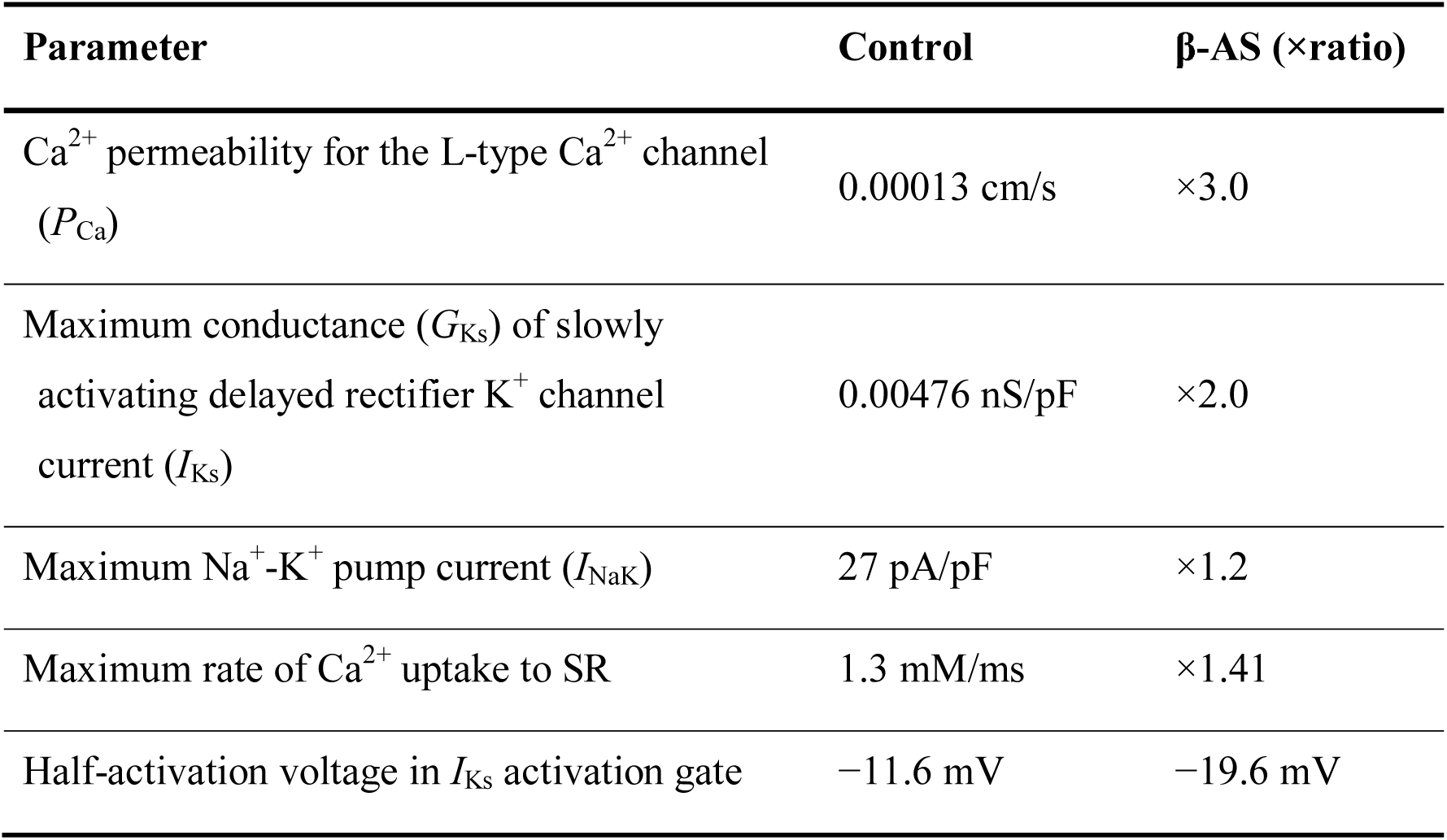
Modification parameters for the condition of *β*-adrenergic stimulation (*β*-AS).

**Table S3.** Comparison with experimental results in ΔSIV mice.

| | | 100%$g_{\text{Na,LM}}$<br>(WT) | 35%$g_{\text{Na,LM}}$<br>($\Delta\text{SIV}$) | |
| --- | --- | --- | --- | --- |
| Peak $I_{\text{Na,LM}}$<br>(pA/pF) | Tsumoto et al | 329.3 | 138.7 | 57.9 % ↓ |
| | Shy et al (10) | $301.1 \pm 25.4$ | $115.0 \pm 11.9$ | $61.8 \pm 4.0$ % ↓ |
| CV (cm/s) | Tsumoto et al | 71.4 | 53.6 | 24.9 % ↓ |
| | Shy et al (10) | $70.1 \pm 4.4$ | $54.2 \pm 2.9$ | $22.7 \pm 0.7$ % ↓ |

Tsumoto et al, Present Study; Shy et al, (10)

